# Extremely distinct microbial communities in closely related leafhopper subfamilies: Typhlocybinae and Eurymelinae (Cicadellidae, Hemiptera)

**DOI:** 10.1101/2024.09.19.613942

**Authors:** Michał Kobiałka, Dariusz Świerczewski, Marcin Walczak, Weronika Urbańczyk

**Author notes:** Address correspondence to Michał Kobiałka.

## Abstract

Among the Hemiptera insects, a widespread way of feeding is sucking sap from host plants. Due to diet poor in nutrients, these insects enter into obligate symbiosis with their microorganisms. However, within the Cicadellidae family, there is a relatively large group of mesophyll feeders – Typhlocybinae that is considered to be devoid of symbiotic companions. In this work, we examine the composition of microorganisms in this subfamily and compare the results with their close relatives – the Eurymelinae subfamily. To study the microbiome, we used high-throughput next-generation sequencing (NGS, Illumina) and advanced microscopic techniques such as transmission electron microscopy (TEM) and fluorescence *in situ* hybridization (FISH) in a confocal microscope. The Typhlocybinae insects have very poor microbial communities in their bodies, these are mainly facultative microorganisms, such as alphaproteobacteria of the genus *Wolbachia* or *Rickettsia*. We detected also the presence of bacteria that can be considered as facultative symbionts e.g. *Spiroplasma, Acidocella, Arsenophonus, Sodalis, Lariskella, Serratia, Cardinium* and *Asaia.* On the other hand, the Eurymelinae group is characterized by a large diversity of the microbial communities, similar to those described in other Cicadomorpha. We find obligate co-symbionts involved in the synthesis of essential amino acids such as *Sulcia,* betaproteobacteria related to genus *Nasuia* or gammaproteobacteria *Sodalis*. In other representatives, we observed symbiotic yeast-like fungi from the family Ophiocordycipitaceae and within some genera we discovered *Arsenophonus* bacteria inhabiting the interior of *Sulcia* bacteria. Additionally, we investigated the transovarial transmission of obligate symbionts, which occurs via infection of the ovaries of females.

**Importance:** The Typhlocybinae and Eurymelinae leafhoppers differ significantly in their symbiotic communities. This is undoubtedly due to their different diets, as Typhlocybinae insects feed on parenchyma, richer in nutrients, while Eurymelinae, like most representatives of Auchenorrhyncha, consume sap from the phloem fibers of plants. Our work presents comprehensive studies of 42 species belonging to two above-mentioned, so far poorly known Cicadomorpha subfamilies. Phylogenetic studies we conducted confirm that the insects from the groups studied have a common ancestor. Since obligate symbionts, having a reduced genome, may affect the reduction of their host’s adaptation to changing environmental conditions, e.g. temperature, and facultative microbiomes may influence the increase in such adaptation and expansion of host niche space. Therefore, Typhlocybinae species may show greater resistance to future climate change than representatives of the Eurymelinae. The research that considers the role of ecological niches in microbiome composition is essential in the era of climate change.

## Introduction

The Typhlocybinae and Eurymelinae are closely related insect subfamilies, belonging to the Cicadellidae, known as leafhoppers – the largest family of Cicadomorpha, which collectively with Fulgoromorpha are placed within Hemiptera (1, 2). Moreover, the Typhlocybinae is the second largest subfamily of the family Cicadellidae (3–5), however recent samplings of tropical faunas indicate that most species of Typhlocybinae remain undescribed, so this taxon can be far larger than any other leafhopper subfamily (4). This monophyletic group of hemipterans includes many plant pests and invasive species that are rapidly spreading across the globe, thus it is claimed that they may be ideal markers of climate change (6–10). The Typhlocybinae group is divided into 7 tribes and even in the European fauna, all currently distinguished tribes of this group have their representatives. The Eurymelinae is the most closely related subfamily to the Typhlocybinae (2). This subfamily is regarded as a paraphyletic group of arboreal leafhoppers, which comprises monophyletic groups, formerly treated as subfamilies, i.a. Eurymelini, Idiocerini and Macropsini (11). The research by Xue and coworkers (11) showed that the Eurymelinae includes 11 tribes, and there are members of two tribes – Idiocerini and Macropsini in Europe.

Although the leafhoppers from the subfamilies mentioned above have a lot in common and often occupy similar ecological niches, however, differ slightly in their feeding mode. The Typhlocybinae are mostly arboreal species, preferentially feeding on the parenchyma of the leaf of their host plants (9, 12) Insect pierces the cuticle and epidermis of the leaf blade using piercing and sucking mouthparts, then sucks up the content of the mesophyll cells. The species belonging to the Eurymelinae are mostly monophagous forms that feed mainly on trees or shrubs. Although they often share a common habitat with Typhlocybinae leafhoppers, they are exclusively phloem feeders. It is commonly believed, that as a rule, leafhoppers feed on plant sap – phloem or xylem from grasses, shrubs and trees, regards to this fact, their diet is supplemented with essential amino acids derived from obligate symbiotic microorganisms (bacteria and/or fungal yeast-like symbionts) (13–16).

Since most hemipterans feed on phloem or xylem sap of plants deficient in essential nutrients, they are host to obligate symbiotic microorganisms: bacteria or fungi (termed yeast-like symbionts). The symbionts of hemipterans are traditionally divided into two groups: obligate symbionts (also termed primary symbionts) and facultative symbionts (known also as secondary symbionts) (13–15). Primary symbionts occur in all the members of particular taxa because they infected the ancestor of this group of insects. Facultative symbionts are younger insect associates, therefore they are often present only in some populations. Obligate symbionts supply their host with nutrients missing from their diet: essential amino acids, cofactors and vitamins (15, 17, 18). Since symbiotic bacteria have strongly reduced genomes, they cannot be cultivated outside the body of the host insect, on laboratory media (19–21). The obligate symbionts are harbored in specialized cells of host insects called bacteriocytes (if bacteria) or mycetocytes (if fungal symbionts), which are grouped into large, paired, distinctively colored structures termed bacteriomes or mycetomes. These microorganisms are transovarially (vertically) transmitted between generations. Facultative symbionts are younger insect associates, therefore very often only some populations may harbor secondary symbionts of different systematic affinity, e.g. *Wolbachia, Rickettsia, Arsenophonus* and *Sodalis*. Facultative symbionts may occur both in bacteriocytes and in other types of insect cells (e.g. in fat body cells) or extracellularly (e.g. in hemolymph). Their function in many cases remains unclear, they may protect the host from heat stress, fungal pathogens and parasitic hymenopterans (22–27).

In contrast to the situation in remaining hemipterans (possess only one type of obligate symbiont), in Cicadomorpha and Fulgoromorpha several types of obligate symbionts are responsible for the synthesis of essential amino acids, therefore they are called "coprimary symbionts" (18, 28–30). Most cicadomorphans and fulgoromorphans are host to ancient symbionts: the bacterium *Candidatus Sulcia muelleri* (phylum Bacteroidetes) and the betaproteobacterial symbiont. During further evolution, in some lineages of Auchenorrhyncha, additional microorganisms have been acquired or the betaproteobacterium has been replaced by other bacteria or yeast-like fungal symbionts (28, 31, 32). Nowadays most cicadomorphans and fulgoromorphans are host to ancient symbionts: bacterium *Sulcia* and the betaproteobacterial symbiont, e.g. *Nasuia deltocephalinicola* in Deltocephalinae leafhoppers;

*Zinderia insecticola* in Clastopteridae spittlebugs; *Vidania fulgoroidea* in Fulgoromorpha planthoppers (29–32).

There are few data on symbiotic associations in members of the Typhlocybinae group. It is hypothesized that these leafhoppers are the only auchenorrhynchans that do not harbor obligate symbionts due to the fact they feed on the parenchyma of leaves (14, 33). So far bacteriomes with bacteria have not been found in representatives of this group. No evidence of transovarial transport of these microorganisms has been reported. Up to date, some facultative bacteria have been detected in bodies of Typhlocybinae insects. Molecular studies indicate that some species are infected with *Wolbachia* or *Ricketssia* and these infections potentially spread among populations by horizontal transmission (9, 34–37).

It is well known that symbiosis plays an important role in the ecology and evolution of both partners, symbiont and host-insect. Many studies prove that the distribution and composition of facultative symbionts may be dependent on biotic and abiotic factors (38, 39). Moreover, symbionts are considered to play a significant role in the diversification and expansion of the ecological niche of their host (40, 41). Results of the recent studies suggest that microbial symbionts may depend on their host response toward abiotic stressors, such as changes in temperature or humidity, by expanding or constraining abiotic niche space (42–44). Because of genome reduction, many obligate symbionts are unable to cope with temperature changes and are considered as their hosts "Achilles’ heels" in the context of temperature stress (43, 45–47). In the era of rapid environmental changes, understanding these relations seems to be crucial for the protection of biodiversity.

In this work, we verified the hypothesis that the Typhlocybinae is a group of leafhoppers that harbors no obligate symbionts and then we examined the symbiotic systems of their relatives – Eurymelinae leafhoppers. We carried out complex research on the symbiotic communities of the members of different tribes: taxonomic composition, occurrence, morphology, localization in the insect body, vertical transmission and phylogeny. By combining morphological and molecular-based approaches (TEM, FISH, and Illumina NovaSeq 6000 based amplicon sequencing), we addressed the following three goals: 1) compare the composition of obligatory and facultative symbiotic communities of 42 species belonging to two studied subfamilies; 2) evaluate whether examined insects exhibit convergent evolution with their symbionts; 3) assess the role of insect phylogeny and diet on composition of symbiotic microorganisms.

## Results

### Eurymelinae symbiotic systems

We examined the composition of the microbiome for the following 18 Eurymelinae species: *Acericerus ribauti, Balcanocerus larvatus, Idiocerus lituratus, Idiocerus stigmaticalis, Macropsis fuscinervis, Macropsis prasina, Macropsis vicina, Oncopsis alni, Oncopsis carpini, Oncopsis flavicollis, Pediopsis tiliae, Populicerus albicans, Populicerus confusus, Populicerus nitidissimus, Populicerus populi, Tremulicerus distinguendus, Tremulicerus tremulae, Viridicerus ustulatus.* The presence of *Sulcia* bacteria was observed in all 66 examined individuals (Fig. 1(I) and (II)). Based on the sequences of these bacteria, we created a cladogram that divides the Eurymelinae leafhoppers into three distinct groups, the first of which is mainly composed of representatives of the genera: *Acericerus, Balcanocerus, Idiocerus* and *Tremulicerus* (tribe Idriocerini); the second: *Populicerus* and *Viridicerus* (tribe Idiocerini); the third: *Macropsis*, *Pediopsis* and *Oncopsis* (tribe Macropsini) (Fig. 1(I)). Betaproteobacteria closely related to *Nasuia* were found in all tested leafhoppers except for the representatives of the genus: *Acericerus, Macropsis* and *Pediopsis*. However, in the body of the mentioned genera, there were symbiotic fungi, *Hirsutella* in *Acericerus* insects and *Ophiocordyceps* in *Macropsis* insects. Alphaproteobacteria of the genera *Wolbachia* and *Rickettsia* are present in almost all examined leafhoppers (Fig. 1(I) and (II)). In some insects, mainly the species of the genus *Tremulicerus* and *I. lituratus*, we demonstrated the presence of *Diploricketssia* bacteria. The *Sodalis* gammaproteobacteria were a relatively common microorganism, with a large number of sequence reads obtained from some representatives of the species *T. tremulae, P. confusus, P. populi, M. prasina, M. fuscinervis, O. alni, O. falvicollis* (Fig. 1(I) and (II)). The genus *Idiocerus* (except *I. lituratus*) and the species *P. confusus* are characterized by the presence of gammaproteobacteria from the genus *Arsenophonus*. In addition to the microorganisms mentioned above, we reported in individual cases, the presence of the following bacteria: *Hepantincola, Spiroplasma, Stenotrophomonas* and *Pseudomonas* (Fig. 1(I) and (II)).

**Fig. 1.**
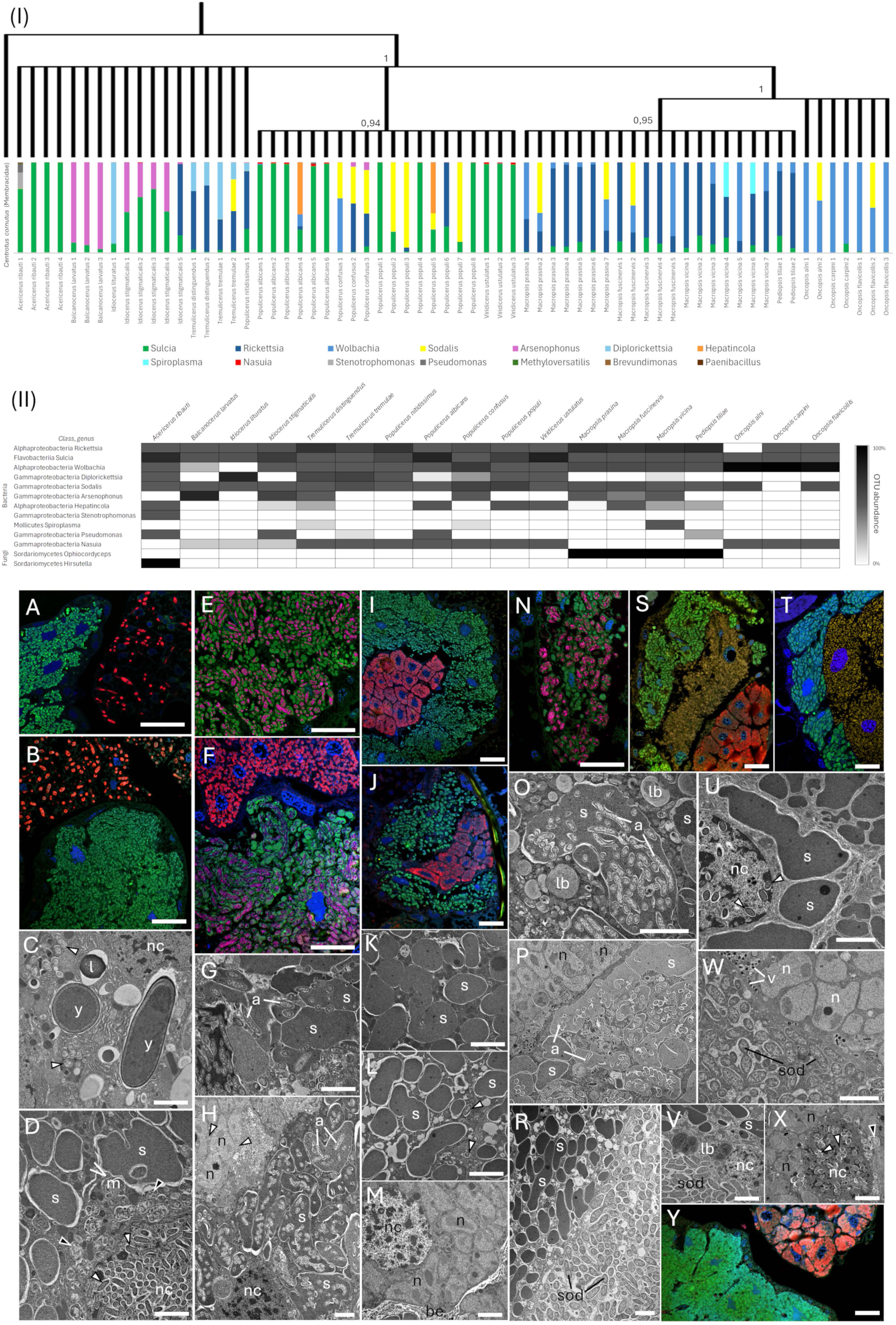
Eurymelinae symbiotic systems. (I) Cladogram based Bayesian analysis of the most abundant sequences of the bacteria *Sulcia* for each individual, showing phylogeny between Eurymelinae species, posterior probabilities are present above the nodes, outgroup *Sulcia* of *Centrotus cornutus* (MN082139.1) (50); below the graphs showing the percentage relative abundance of bacteria for each sample based on the number of sequences reads. (II) OTUs relative abundance of both bacterial and fungal microorganisms, collectively for each Eurymelinae species. (A) *A. ribauti*, bacteriome containing *Sulcia* bacteria (green), surrounded by a fat body where yeast-like symbionts of the *Hirsutella* genus are present (red). (B) *M. vicina,* bacteriome containing *Sulcia* bacteria (green), surrounded by a fat body where yeast-like symbionts of the *Ophiocordyceps* genus are present. (C) *M. prasina, Ophiocordyceps* yeast-like symbionts (y) in insect fat body. (D) *P. tilliae,* bacteria *Sulcia* (s) in bacteriocyte, note bacteria *Wolbachia* (white arrowheads) inside cell nucleus (nc). (E and F) *I. stigmaticalis, Arsenoponus* bacteria (purple) resident inside *Sulcia* bacteria (green), bacteria *Nasuia* (red) present in bacteriocytes forming a separate zone. (G) *B. larvatus, Arsenoponus* bacteria (a) infecting bacteria *Sulcia* (s). (H) *I. stigmaticalis,* bacteriocytes filled with bacteria *Sulcia* (s) infected by bacteria *Arsenophonus* (a), in the upper left corner a fragment of a bacteriocyte occupied by bacteria *Nasuia* (n). (I) *P. albicans,* (J) *P. populi,* bacteriome composed of two zones, bacteriocytes filled with *Sulcia* bacteria (green) and *Nasuia* bacteria (red). (K) *P. albicans,* (L) *P. populi,* pleomorphic *Sulcia* bacteria inhabiting bacteriocytes. (M) *P. albicans,* the fragment of a bacteriocyte filled with pleomorphic *Nasuia* bacteria (n). (N) *P. confusus,* bacteriocytes filled with *Sulcia* bacteria (green) infected by *Arsenophonus* bacteria (purple). (O and P) *P. confusus,* bacteria *Sulcia* (s) filled with many rod-shaped *Arsenophonus* bacteria (a), in the upper left corner the fragment of a bacteriocyte occupied by bacteria *Nasuia* (n). (R) *P. nitidissimus,* two zones of a bacteriome, on the left pleomorphic *Sulcia* bacteria (s), on the right rod-shaped *Sodalis* bacteria (sod). (S) *T. tremulae,* (T) *V. ustulatus,* bacteriome where bacteriocytes filled with various bacteria form three distinct zones: *Sulcia* (green), *Sodalis* (yellow) and *Nasuia* (red). (U) *T. distinguendus,* pleomorphic *Sulcia* bacteria (s) inhabiting bacteriocytes, note rod-shaped alphaproteobacteria present in the cell nucleus (white arrowheads). (W) *T. distinguendus,* two zones of bacteriome, on the left rod-shaped *Sodalis* bacteria (sod), on the right pleomorphic *Nasuia* bacteria (n), note the viruses (v) resident in bacteriome epithelium between zones. (V) *V. ustulatus,* two zones of bacteriome, on the left pleomorphic *Sulcia* bacteria (s), on the right rod-shaped *Sodalis* bacteria (sod). (X) *O. alni,* pleomorphic *Nasuia* bacteria (n) inhabiting bacteriocytes, note rod-shaped alphaproteobacterial present in the cell nucleus (white arrowheads). (Y) *O. flavicollis,* bacteriocytes filled with *Sulcia* (green) and *Nasuia* (red) bacteria forming two separate bacteriomes. (A, B, E, F, I, J, N, S, T, Y) Confocal microscope; cell nuclei, DAPI staining (blue); scale bar – 40 µm. (C, D, G, H, K-M, O-R, U-X) TEM; be, bacteriome epithelium; black arrowheads, rod-shaped gammaproteobacterium; l, lipid droplet; lb, lamellar body; m, mitochondrion; nc, cell nucleus; white arrowhead, rod-shaped alphaproteobacterium; scale bar – 3 µm.

During microscopic analysis, we observed the presence of specialized organs, called bacteriomes, composed of mostly spherical cells, called bacteriocytes, which are the habitat of obligate symbionts (Fig. 1A, B, E, F, I, J, N, S, T, Y; Mov. 1, 2, 3). Bacteriomes occur as paired structures, two pairs in number, on the sides of the body, in the upper part of the abdomen, near the reproductive organs. They are surrounded by a single-layered epithelium, sometimes called the bacteriome sheath. In the bodies of the representatives of the genus: *Acericerus, Macropsis* and *Pediopsis* we noted the occurrence of bacteriomes occupied only by the bacteria *Sulcia*, additionally, yeast-like microorganisms were present in the insect’s fat body (Fig. 1A-D, Mov. 1). In other leafhoppers, the bacteriomes formed zones, each surrounded by its epithelium, composed of bacteriocytes, whose cytoplasm was filled with specific types of bacteria: *Sulcia* and *Nasuia*. Sulcia bacteriocytes formed the outer zone of bacteria, while *Nasuia* bacteriocytes filled the central part (Fig. 1E, F, I, J; Mov. 2). In the members of the genus *Oncopsis*, bacteriocytes of *Sulcia* and *Nasuia* formed separate bacteriomes (Fig. 1Y). Bacteria of the genus *Sulcia* were quite large microorganisms (10-12 µm in length), they were pleomorphic, sometimes slightly elongated, electron-dense, and therefore darker in electron microscopy images in contrast to *Nasuia* bacteria (Fig. 1D, G, K, L, R, U). Betaproteobacteria related to the genus *Nasuia* also had a pleomorphic shape, but unlike *Sulcia* bacteria, they were more rounded (Fig. 1H, W, X), sometimes branching (Fig. 1M and P), and were electron-transparent in TEM images.

In individuals of the species *B. larvatus*, *I. stigmaticalis* and *P. confusus*, we observed an unusual symbiotic system, the gammaproteobacteria *Arsenophonus* were present inside bacteria of the genus *Sulcia* (Fig. 1E-H, N-P; Mov. 3). Rod-shaped, strongly elongated *Arsenophonus* bacteria did not settle each of all *Sulcia* bacteria, and some of them occurred in the cytoplasm of *Sulcia* bacteriocytes (Fig. 1G, H, O, P). These free bacteria were often removed by lysosomes. This is why we observed in some places the forming of lamellar bodies as a result of the destruction of bacteria recognized as pathogens (Fig. 1O). In turn, rod-shaped gammaproteobacteria *Sodalis* formed a separate zone, between *Sulcia* and *Nasuia,* within the bacteriomes of their hosts: *P. nitidissimus, T. tremulae, T. distinguendus* or *V. ustulatus* (Fig. 1R, T, W, V). We sometimes observed *Sodalis* bacteria and viruses in the epithelium separating the individual zones, lamellar bodies were visible in the places where they were destroyed (Fig. 1W and V).

Representatives of alphaproteobacteria such as *Wolbachia* and *Rickettsia* occurred in many locations in the bodies of the studied insects. *Wolbachia* bacteria were present inside the bacteriomes (Fig. 1D, H, L, U, X, 3A-C) but also in the fat body (Fig. 1C and 3P), midgut epithelium (Fig. 3G, J, L) and reproductive organs - tropharia in the ovaries of females (Fig. 3O and R). Bacteria *Wolbachia* and *Rickettsia* were indistinguishable in TEM images due to their very similar morphology: small, rod-shaped, slightly elongated, and electron-transparent. It is worth mentioning, that bacteria of the *Wolbachia* genus were quite often found inside the cell nuclei of various cell types: bacteriocytes both with *Sulcia* and *Nasuia* (Fig. 1D, U, X, 3C), midgut epithelium (Fig. 3J), fat body (Fig. 3P) and trophocytes (Fig. 3R). We assumed that above mentioned bacteria were *Wolbachia* because this is the genus that has been previously described inside cell nuclei (48). Additionally, apart from alphaproteobacterial, we observed yeast-like fungal symbionts in the cytoplasm of the midgut epithelium of *M. vicina* insects (Fig. 3L).

### Typhlocybinae symbiotic systems

We examined the composition of the microbiome for the following 24 Typhlocybinae species: *Alebra albostriella, Chlorita paolii, Edwardsiana lethierryi, Emelyanoviana mollicula, Empoasca pteridis, Erythria aureola, Eupterycyba jucunda, Eupteryx aurata, Eupteryx calcarata, Eupteryx cyclops, Eupteryx notata, Eurhadina pulchella, Fagocyba cruenta, Forcipata citrinella, Kybos populi, Kybos virgator, Linnavuoriana sexmaculata, Notus flavipennis, Ribautiana tenerrima, Typhlocyba quercus, Zygina hyperici, Zygina rubrovittata, Zyginella pulchra, Zyginidia pullula. Wolbachia* bacteria were present in almost all species, except *E. lethierryi, E. cyclops* and *Z. pulchra*. The second most common bacteria was *Rickettsia* (Fig. 2A). Alphaproteobacteria were observed mainly in the insect midgut epithelium (Fig. 3D-F, K) and fat body (Fig. 3M and N). *E. cyclops* and *K. populi* were infected with *Spiroplasma* bacteria. We detected bacteria of the genus *Acidocella* in higher amounts in individuals of the species: *E. mollicula*, *Z. pulchra* and *Z. pullula*. We did not detect the presence of obligate symbionts in the samples. However, we have noted the occurrence of some facultative symbionts*: Arsenophonus, Sodalis, Lariskella, Serratia, Cardinium* and *Asaia* (Fig. 2A). The samples also included bacteria from chitinous cuticles or the guts of insects.

**Fig. 2.**
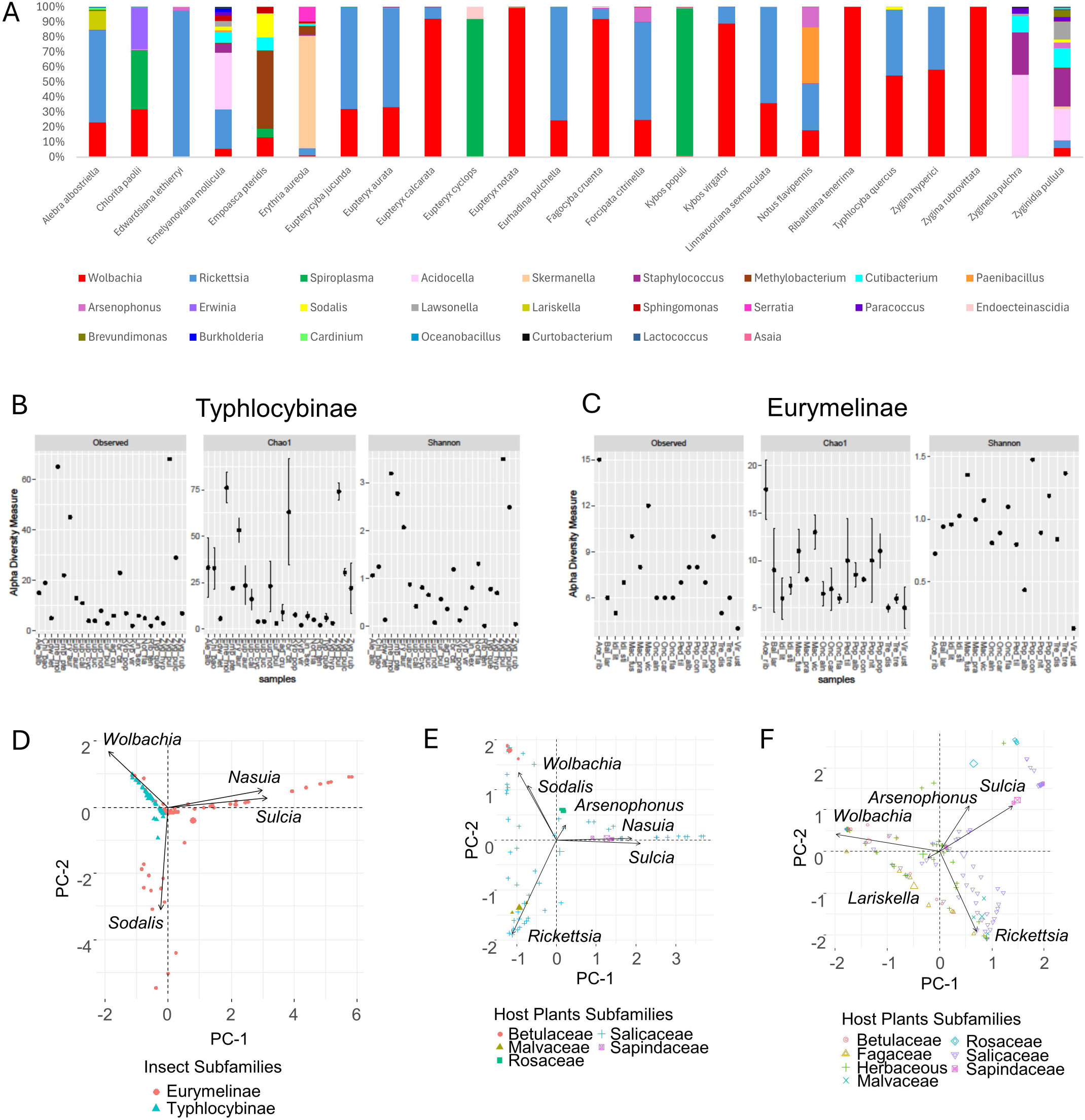
Typhlocybinae and Eurymelinae microbiomes comparison. (A) The percentage relative abundance of bacteria for Typhlocybinae species based on the number of sequence reads. (B) Typhlocybinae, (B) Eurymelinae, alpha diversity indices given for each species (species name abbreviations are given in Table S1). (C) The PCA analysis, Typhlocybinae and Eurymelinae microbiome diversity, the arrows indicate the loadings - the most important types of bacteria, differentiating two groups of insects. (D) The PCA analysis, Eurymelinae microbiome diversity for various plant subfamilies, which insects feed on, the arrows indicate the loadings - the most important types of bacteria, differentiating groups of host plants. (E) The PCA analysis, Typhlocybinae and Eurymelinae microbiome diversity for various plant subfamilies, which insects feed on, the arrows indicate the loadings - the most important types of bacteria, differentiating groups of host plants.

**Fig. 3.**
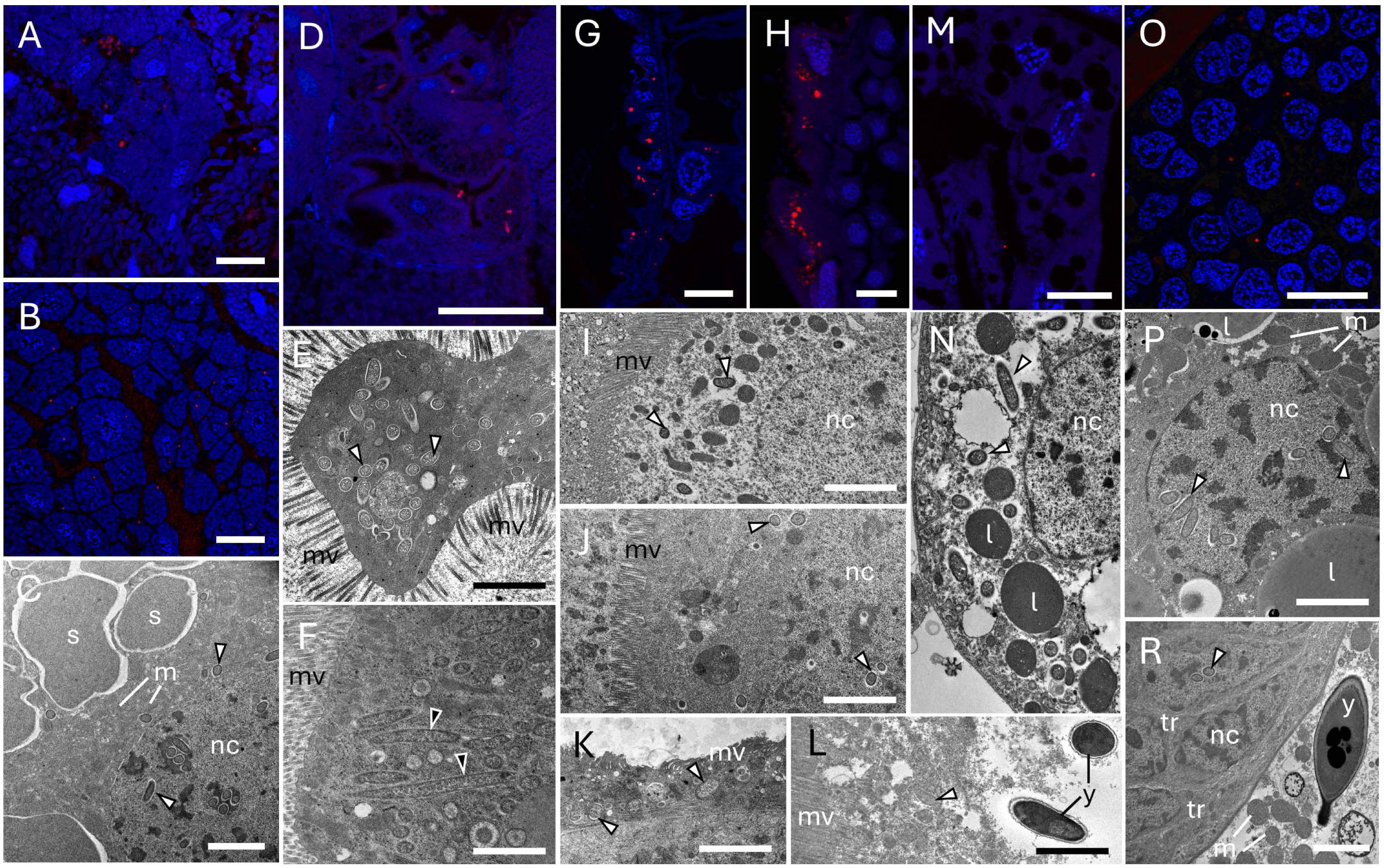
Facultative bacteria in various tissues of Typhlocybinae and Eurymelinae insects. (A) *P. populi,* (B) *P. albicans, Wolbachia* bacteria (red) present in *Nasuia* bacteriocytes zone. (C) *V. ustulatus*, alphaproteobacteria (white arrowheads) occurring in cytoplasm and nucleus (nc) of *Sulcia* (s) bacteriocyte. (D) *E. aurata, Wolbachia* bacteria (red) present in midgut epithelium. (E) *F. citrinella,* alphaproteobacteria (white arrowheads) settling the cell of the midgut epithelium. (F) *E. lethierryi,* gammaproteobacteria (black arrowheads) present in midgut epithelium. (G) *P. albicans,* (H) *V. ustulatus, Wolbachia* bacteria (red) occurring in midgut epithelium. (I) *Z. hyperici* (J) *V. ustulatus* (K) *T. quercus* (L) *M. vicina,* alphaproteobacteria (white arrowheads) present in midgut epithelium. (M) *E. calcarata, Wolbachia* bacteria (red) in insect fat body. (N) *F. citrinella,* alphaproteobacteria (white arrowheads) settling fat body cells. (O) *P. albicans, Wolbachia* bacteria (red) occurring in trophocytes (ovary). (P) *T. tremulae,* alphaproteobacteria (white arrowheads) settling nucleus in fat body. (R) *P. tiliae,* alphaproteobacteria (white arrowheads) in trophocytes (tr) (ovary), note yeast-like microorganism (y) in the fat body, lower right corner. (A, B, D, G, H, M, O) Confocal microscope; cell nuclei, DAPI staining (blue); scale bar – 40 µm. (C, E, F, I-L, N, P, R) TEM; l, lipid droplet; m, mitochondrion; mv, intestinal microvilli; nc, cell nucleus; y, yeast-like microorganism; scale bar – 3 µm.

### Phylogenetic results

The phylogeny of Eurymelinae based on sequences of the bacteria *Sulcia* is shown in Fig. 1(I) and described at the beginning of this chapter. Unfortunately, the lack of obligate symbionts in individuals of Typhlocybinae has prevented this procedure from being performed in this group. However, based on the sequence of the insect COI (cytochrome C oxidase I) gene, we have constructed a cladogram showing the relationships between all studied leafhoppers (Fig. 4). Representatives of the Macropsini tribe constitute separate group from other leafhoppers. The remaining members of Eurymelinae form a separate clade within Typhlocybinae leafhoppers, showing closest relationships to members of the genera *Alebra, Chlorita, Empoasca* and *Kybos*. The remaining genera of Typhlocybinae form successive clades among themselves, within tribes and genera. The most outlying group of them is formed by representatives of the species *Z. pulchra*.

**Fig. 4.**
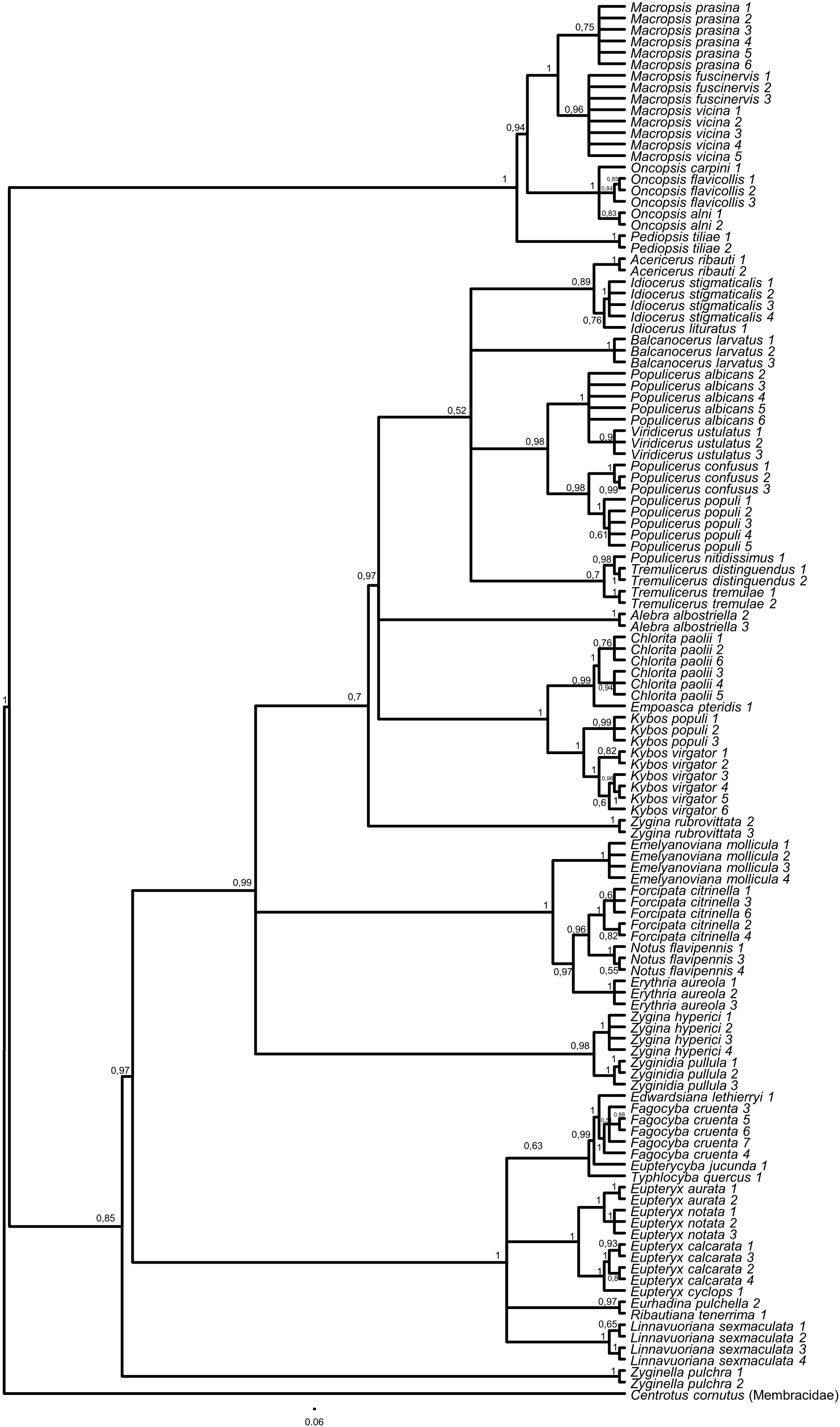
Cladogram based Bayesian analysis of COI insect gene, showing phylogeny between both Typhlocybinae and Eurymelinae species; posterior probabilities are present above the nodes; outgroup *C. cornutus* (MW536003.1) (88).

#### Transovarial transmission of microorganisms

During microscopic observations, we observed transovarial transmission of symbiotic microorganisms only in representatives of the Eurymelinae subfamily. Symbiotic bacteria migrate from the bacteriome towards the posterior pole of the terminal oocyte, located at the end of the ovariole - the basic component of the ovary. If we observed symbiotic fungi in species, we noted their migration from the fat body to the oocytes (Fig. 5A). The symbionts infect follicular cells, filling their cytoplasm (Fig. 5A-C, E). During this process, *Sulcia* and *Nasuia* bacteria change their shape from pleomorphic (still in Fig.5E) to more spherical (Fig. 5B, C, F). We have not noted this phenomenon for symbiotic fungi (Fig. 5F). Then the microorganisms enter the space between the oocyte and the follicular cells, called the perivitelline space. At this point, symbiotic microorganisms form a structure called a "symbiont ball", which in this case has a "cap-like" shape (Fig. 5B, D; Mov. 4, 5). Within the "symbiont ball", we observed symbionts that were present in the host bacteriome, e.g. *Sulcia, Nasuia, Sodalis* for *V. ustulatus* (Fig. 5G), or *Sulcia, Arsenophonus, Nasuia* for *B. larvatus* – *Arsenophonus* bacteria were present as well as in bacteriome inside the *Sulcia* bacteria (Fig. 5H).

**Fig. 5.**
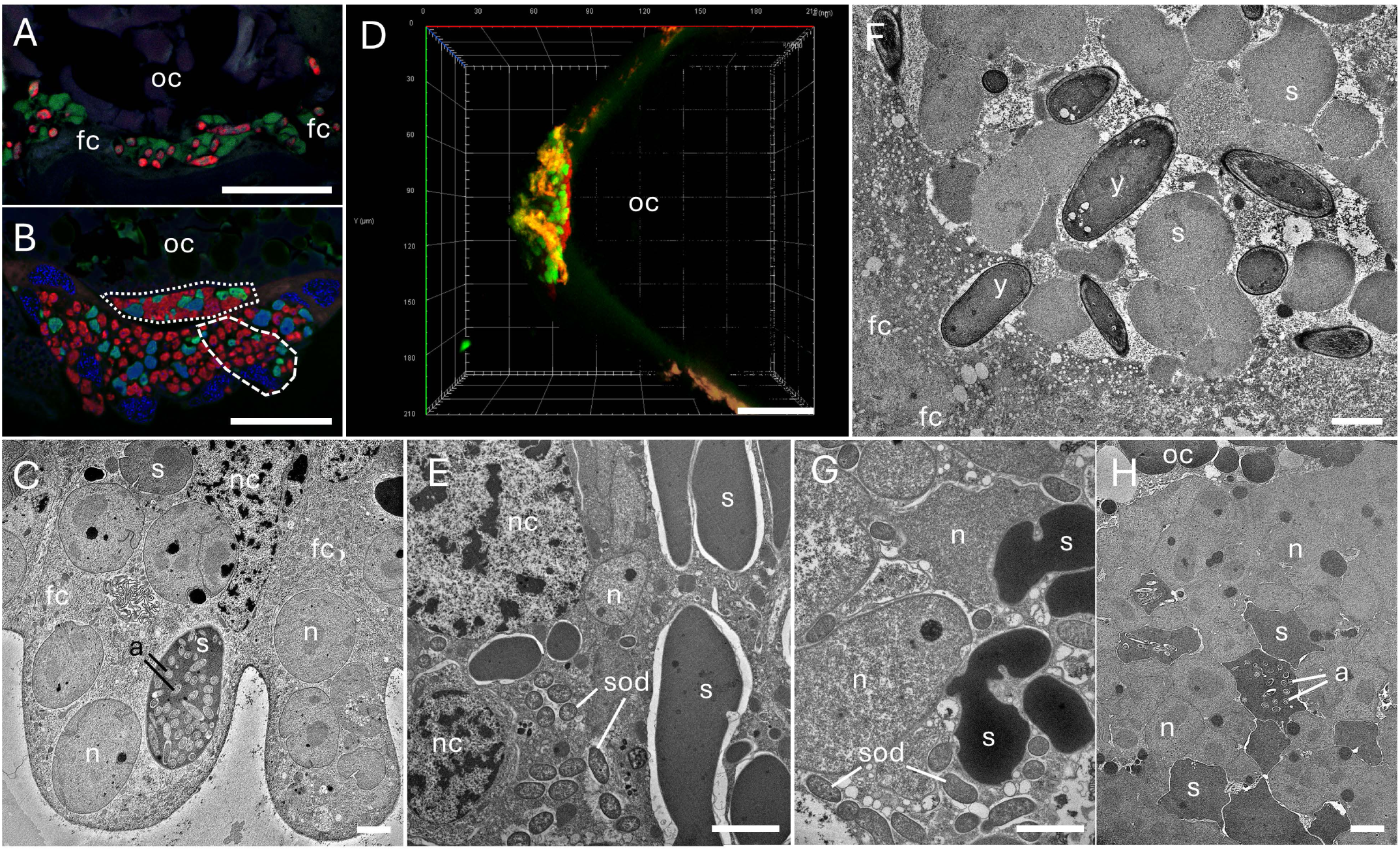
Transovarial transmission of bacteria in Eurymelinae leafhoppers. (A) *M. vicina, Sulcia* bacteria (green) and *Ophiocrodyceps* yeast-like symbionts (red) infecting follicular cells (fc) surrounding the posterior pole of the oocyte (oc). (B) *P. albicans, Sulcia* (green) and *Nasuia* (red) bacteria infecting follicular cells (one of them marked by a dashed line) and forming cap-like "symbiont ball" (dotted line) in invagination of oolemma. (C) *B. larvatus, the* fragment of follicular cells (fc) filled with *Sulcia* (s) and *Nasuia* (n) bacteria, note *Arsenophonus* bacteria (a) inside *Sulcia* (s). (D) *P. albicans,* 3D view of "symbiont ball" consisted of *Sulcia* (green) and *Nasuia* (red), note the faint green and red flashes of autofluorescence on the chorion. (E) *T. tremulae,* three types of bacteria: *Sulcia* (s), *Sodalis* (sod) and *Nasuia* (n) present in follicular cells. (F) *M. prasina,* the fragment of *"symbiont ball"* consisted of *Sulcia* bacteria (s) and *Ophiocordyceps* yeast-like symbionts (y), note follicular cell (fc) in the bottom left corner. (G) *V. ustulatus, the* fragment of "symbiont ball" consisted of *Sulcia* (s), *Sodalis* (sod) and *Nasuia* (n) bacteria. (H) *B. larvatus,* the fragment of "symbiont ball" consisted of *Sulcia* (s) and *Nasuia* (n) bacteria, note *Arsenophonus* bacteria (a) inside *Sulcia* (s), note the fragment of an oocyte (oc) in the upper left corner. (A, B, D) Confocal microscope; cell nuclei, DAPI staining (blue); oc, oocyte; scale bar – 40 µm. (C, E, F-H) TEM, nc, follicular cell nucleus; scale bar – 3 µm.

## Discussion

### Microbiome comparison

Despite their clear phylogenetic relationship, the leafhoppers from the subfamilies Typhlocybinae and Eurymelinae differ in their microbiome (Fig. 1(I) and 2A). We noticed that most of the Eurymelinae individuals showed higher microbial diversity compared to Typhlocybinae (Fig. 2B and C -alpha diversity, Chao and Shannon index). We confirm that Typhlocybinae is a group of leafhoppers devoid of obligate symbiotic microorganisms and with a much poorer microbiome diversity. The studied Eurymelinae insects are monophagous, living in trees, and feeding on xylem sap, while the Typhlocybinae suck up the mesophyll contents, having their host plants and trophic specialization more diverse (Table S1). That is why the variability of their microorganisms largely comes down to the diversity of bacteria that make up the guts microflora. Nevertheless, some of the examined species of both subfamilies feed on the same host plant, e.g. host-plant *Carpinus betulus* inhabiting leafhoppers *O. carpini* versus *F. cruenta*; host*-*plant *Salix alba* inhabiting leafhoppers *I. stigmaticalis* versus *E. pteridis, K. virgator*; host-plant *Salix cinerea* inhabiting leafhoppers *M. prasina, P. confusus* versus *L. sexmaculata.* However, in these cases, we noticed only common facultative alphaproteobacteria *Wolbachia* or *Rickettsia*, which are presumably transmitted horizontally (Fig. 1(I) and 2A).

We should point out, that the Eurymelinae microbiome is mainly composed of obligate co-symbionts, responsible for the synthesis of essential amino acids. This is a similar situation to other members of the Cicadellidae family, mainly the subfamily Deltocephalinae (48–61). The main co-symbionts of Eurymelinae are *Sulcia* and betaproteobacteria related to the genus *Nasuia*. There are also two main bacteria that, together with *Sodalis* and *Wolbachia*, distinguish the Eurymelinae microbiome from Typhlocybinae (Fig. 2D).

Phylogenetic position along with diet may influence the host microbiome (62). The occurrence of some bacteria in the insects Typhlocybinae and Eurymelinae was dependent on the host plant. In Eurymelinae group, the main bacteria influencing the grouping of leafhoppers according to their host plants are: *Sulcia, Nasuia, Arsenophonus, Sodalis, Rickettsia* and *Wolbachia* (Fig. 2E). When analyzing two subfamilies together, we observed such trends as the presence of *Rickettsia* bacteria was higher in the insects living on Salicaceae family trees, *Wolbachia* in leafhoppers on herbaceous plants, *Arsenophonus* in insects on Rosaceae plants, and *Lariskella* in leafhoppers on Fagaceae plants (Fig. 2F).

Within the Eurymelinae group itself, symbiotic system diversity can be observed. This is influenced not only by the above-mentioned bacteria *Sulcia, Nasuia, Sodalis* (Fig. 2D) but also by symbiotic fungi and *Arsenophonus* bacteria (Fig. 1(I) and (II)). The Macropsini tribe is a distinctly different group - the content of *Sulcia* bacteria is lower (Fig. 1(I)), the bacteriomes are smaller (Mov. 2), the symbiotic system is supplemented with *Ophiocordyceps* yeast-like microorganisms, occasionally with *Sodalis* bacteria, and individuals from the *Oncopsis* genus do indeed have betaproteobacteria *Nasuia*, but they inhabit separate bacteriomes (Fig. 1Y). The presence of fungi as symbiotic partners of leafhoppers has been observed previously both in Cicadomorpha and Fulgoromorpha (55, 63, 64). In this group, we noted alphaproteobacteria often appearing in large quantities, in individuals from the *Macropsis* genus it was more often *Wolbachia*, and in individuals from the *Oncopsis* genus – *Rickettsia* (Fig. 1(I)). It should be added that these insects differ in their diet, *Oncopsis* insects feed on the sap of trees from the Betulaceae subfamily, while *Macropsis* – on Salicaceae. The remaining representatives classified as the tribe Idiocerini, can be divided into two groups due to their microbiomes. The first, more uniform, phylogenetically coherent group consists of the genera *Populicerus* and *Viridicerus* (middle clade Fig. 1(I)). These leafhoppers have a symbiotic system consisting of *Sulcia* + *Nasuia*, supplemented in some cases by *Sodalis* bacteria. The exceptions here are individuals of the species *P. confusus* – their systems have numerous alphaproteobacteria and *Arsenophunus* bacteria inside *Sulcia* associates. These insects live on plants of the *Salix cinerea* species, unlike the others from this group that feed on trees of the *Populus* genus. The *Arsenophonus* bacterium is considered to be a symbiont that, under stressful conditions or with a smaller amount of food, displaces obligate symbionts, replacing them in the synthesis of amino acids (65). It may also be responsible for the synthesis of vitamin B (66). The occurrence of bacteria in bacteria has already been described (56, 58, 67). The last group of Eurymelinae, distinguished by us due to the microbiome, contains diverse systems, where in addition to *Sulcia* bacteria can be found: 1) symbiotic *Hirsutella* fungi in individuals of the genus *Acericerus* (the only insects among those studied feeding on plants of the *Acer* genus); 2) *Arsenophonus* bacteria inside *Sulcia* bacteria in individuals of the genera *B. larvatus* and *I. stigmaticalis*; 3) *Diploricketssia* bacteria in representatives of the genus *Tremulicerus* and species *I. lituratus* (Fig. 1(I) and (II)). Some *Diploricketssia* bacteria have been described as human pathogens – microorganisms occurring in the genus *Ixodes* (68, 69). *Tremulicerus* insects are also characterized by having significant amounts of *Rickettsia* bacteria in their bodies. We noticed that two *Macropsis* insects were infected with *Spiroplasma* bacteria (greater number of reads). *Spiroplasma* is a microorganism that can be considered as a pathogen, commensal bacteria or even mutualistic partner, their influence has been compared to the *Wolbachia* bacteria (70, 71). Also two individuals, but from the genus *Populicerus*, were infected with the bacterium *Hepanticola*, an extracellular midgut bacterium found in terrestrial isopods, that feeds on the nutrients of its host (72). In some cases, the sequencing results obtained by us revealed bacteria, probably collected from the surface of the insect cuticle or occurring in their digestive tract, e.g. *Stenotrophomonas, Pseudomonas, Brevundimonas*. We do not rule out that the individual infections described above may originate from parasites or parasitoids inhabiting the insect’s body, although our samples were checked for this (during preparation, COI gene sequencing).

In contrast to Eurymelinae, the body of insects from the subfamily Typhlocybinae is inhabited mainly by alphaproteobacterial microorganisms of the genera *Wolbachia* and *Rickettsia*. *Wolbachia* and *Rickettsia* are bacteria often classified as intracellular commensals or pathogens. They inhabit various types of host cells, quite frequently occupying the host bacteriomes or gonads, which may provide greater shelter by removing them from the body and potential vertical transport. It persists in the population and is passed on to the greatest extent by females to offspring. *Wolbachia* bacteria are known to interfere with sex determination in hosts, cause feminization and death of male offspring (73). *Rickettsia* bacteria are most often transmitted horizontally, by parasites, and also inhabit places such as the ovaries, mouthparts, or digestive tract. This microorganism has a positive effect on fertility, survival rate, or accelerated development, but also on the feminization of offspring (74).

In some Typhlocybinae species, we observed the presence of bacteria that can be considered as facultative symbionts e.g. *Spiroplasma, Acidocella, Arsenophonus, Sodalis, Lariskella, Serratia, Cardinium* and *Asaia.* Representatives of the species *Ch. paolii*, *E. pteridis*, and especially *E. cyclops* and *K. virgator* possess *Spiroplasma* bacteria in their bodies. We obtained a relatively large number of *Acidocella* reads from samples originating from *E. mollicula*, *Z. pulchra* and *Z. pullula*. These bacteria were described as occurring in the guts of bees (75), what is more, they belong to the Acetobacteriaceae, a group of bacteria among which bacterial symbionts such as *Asaia* can be found, bacteria distributed in the guts, fatty bodies, and salivary glands, involved in host’s nutrient supply (76). Also, *Arsenophonus* bacteria were detected in some samples and in higher read numbers in *F. citrinella* and *N. flavipennis* individuals from the same habitat, *Carex* plants, suggesting a possibility of horizontal transport. We have detected *Sodalis* bacteria among *E. mollicula, E. pteridis* and *T. quercus.* These microorganisms, as we know, are described not only as facultative but also as obligate symbionts involved in the nutrition of the host, associated with the bacteriome (77). However, we did not observe any bacteriomes during a histological examination of Typhlocybinae. All the tested individuals of the species *A. albostriella* harbor bacteria of the genus *Lariskella* in their bodies. It is an *Ixodes* symbiont, the quantity of which varies mainly depending on the sex of the individuals (78). *Serratia* bacteria occurred in the species *E. aureola* and *Cardninium* endosymbiont in *Z. pullula*, both being described as facultative symbionts of Cicadomorpha (50, 54). The remaining detected microorganisms, often observed in single samples, were probably collected from the insect’s cuticle surface or present in its food, in the host guts.

### Phylogenetic remarks

Based on the obtained cladogram, we can state that the tribe Macropsini is a monophyletic group that separates from the other studied leafhoppers (Fig. 4). Their symbiotic system also confirms this fact. Our studies are consistent with the distinctiveness of this group, shown in earlier works (2, 11). The remaining representatives of Eurymelinae constitute a compact clad more related to the studied Typhlocybinae. This phylogeny of Macropsini and Idiocerini coincides with that shown in the supplementary phylogenetic tree in the work of Dietrich and coworkers (2), where Macropsini is a distinct group related to the Deltocephalinae. In the past, these leafhoppers were considered a separate subfamily. Insects from the tribe Idiocerini are probably a paraphyletic group as indicated in the work mentioned above (2)and confirmed by our studies. Additionally, the diversity of this group can be seen in their symbiotic communities. In our cladogram, the insects of the tribe Idiocerini are most closely related to the genera *Alebra* (tribe Alebrini) and the genera belonging to the tribe Empoascini (genus *Chlorita, Empoasca, Kybos*) (Fig. 4). Alebrini is considered to be phylogenetically the oldest group of Typhlocybinae, from which the remaining representatives of this group may have originated (79). To all of the above-mentioned leafhoppers, the sister groups are Erythroneurini (genus *Zygina, Zyginidia)* and Dikraneurini tribe (genus *Emelyanoviana, Forcipata, Notus, Erythria*). The lower part of the cladogram is filled by a large group of representatives of the Typhlocybini tribe and a small group consistent with the genus *Zyginella* (tribe Zyginellini). To summarize, in the cladogram created in our study, confirms the phylogenetic classification of insects within Typhlocybinae (80).

### Microorganisms transmission

The transmission of microorganisms can occur horizontally or vertically, between generations. Horizontal transmission usually concerns facultative bacteria, mainly representatives of alphaproteobacteria, e.g. *Wolbachia* or *Rickettsia*. It probably occurs between populations or species of the studied Eurymelinae and Typhlocybinae insects, especially when they occur in the same location, on the same host plant. More advanced vertical transmission is performed through infections of ovarioles (components of the female reproductive system), which mainly affects obligate symbionts. We did not observe any traces of transovarial transport in representatives of Typhlocybinae. However, we have observed this kind of transmission in representatives of Eurymelinae, and it occurs according to a mechanism similar to that of other representatives of leafhoppers or even Auchenorrhyncha (48, 50, 58, 81). Microorganisms pass through the follicular epithelium into the perivitelline space, where they form a cap-like structure called a "symbiont ball". This phenomenon confirms the long-term coevolution of these insects with their symbionts.

## Conclusions

In this work, based on comprehensive molecular and microscopic studies, we compare symbiotic systems of two closely related Cicadellidae subfamilies (Typhlocybinae and Eurymelinae). We confirm the hypothesis of the lack of obligate symbionts in the leafhoppers from the subfamily Typhlocybinae. Since representatives of the tribe Idiocerini (Eurymelinae) are closely related to the ancestral Typhlocybinae tribe Alebrini, it is possible that two examined subfamilies have a common ancestor. While Eurymelinae is a subfamily characterized by a large diversity of symbiotic systems (including *Sulcia, Nasuia, Arsenophonus, Sodalis* bacteria, yeast-like fungal symbionts), Typhlocybinae members have a poor microbiome, characterized mainly by the presence of alphaproteobacteria, e.g. *Wolbachia*, which are common in the world of insects and even arthropods (82). It remains enigmatic, what was the reason for the interruption of the close mutual symbiosis and long coevolution that connected these leafhoppers with symbiotic associates. In our opinion, the cause of this phenomenon is the reduction of the genome of obligate symbionts, leading to the deficit in the adaptation of insects to changing environmental conditions, primarily changes in temperature (40, 43, 45). For the same reason, species belonging to Typhlocybinae may show greater resistance to future climate change than members of Eurymelinae. Studied insects differ in their feeding habits, Eurymelinae feeds on phloem sap, and Typhlocybinae feeds on mesophyll tissue. Additionally, unlike the monophagous Eurymelinae, in the Typhlocybinae group can be found insects of different trophic specialties, feeding on various trees, shrubs or herbaceous plants. It is possible that the lack of dietary specialization (and therefore a wide dietary niche) also results from the lack of primary endosymbionts in Typhlocybinae. We can also notice a particular relationship between the composition of microbiome and the main host plant, with a large share of *Rickettsia* in leafhoppers feeding on trees belonging to the Salicaceae family. Typhlocybinae is a reasonably large group, found in many habitats on our planet, especially abundant in a tropical zone. This is why we intend to undertake future research on the understanding of the symbiotic systems of this group collecting individuals from many areas and to compare the results with the data referring to the ecological niches they inhabit.

## Material and methods

### Molecular identification

In this study, we examined 158 individuals belonging to 42 species (18 of Eurymelinae and 24 of Typhlocybinae), taken from several localities in southern Poland. Tables with collected specimens, collection localities, as well as information about species feeding specificity and host plant are provided in the appendix (Table.S1). The adult specimens, destined for the molecular analysis, were fixed in 100% ethanol. Genomic DNA was extracted from specimens using the commercially available kits for DNA isolation (GeneMATRIX Bio-Trace DNA Purification Kit EURx) following manufacturer protocol and subsequently stored at 4 °C for further analysis. Symbiotic microorganisms associated with examined species were determined based on their 16S rDNA (bacteria) and ITS1, ITS2 (fungal symbionts) sequences which are commonly used in taxonomic studies (83, 84). As the bacterial 16S rRNA gene contains highly conservative and hypervariable regions (V1-V9), the analysis of its sequences (especially variable region V3-V4) let us determine the systematic affiliation of bacteria and the relationships among them (85). The quantity of DNA was estimated with Qubit by measuring using fluorescence-based assays. Genomic DNA isolation was confirmed by amplifying the 16S rDNA and 18S rDNA using a pair of primers: V3-V4: F357, R805 and NS1, FS2 (Table S2) and visualized using electrophoresis on agarose gel. The samples were sequenced by SEQme s.r.o. (Czech Republic) using the Illumina NovaSeq 6000 SF Flow Cell. The sequencing libraries were prepared in the Nextera Technology kit using primers presented in Table S2. All reads were subjected to quality verification using FastQC and MultiQC. The reads were demultiplexed, forward and reverse sequences were joined using Fastq-join, then filtered, poor quality adapter sequences and chimeras were removed. The reads were classified into operational taxonomic units (OTUs) using Usearch and using the clustering method at a similarity threshold of 97% sequence identity. Then, the OTUs were assigned to the genus level based on databases (Silva 138.1, Unite 9, NCBI Genbank). Indices for alpha diversity we obtained using the Phyloseq package in R Studio. To perform the PCA analysis and create plots we used the Factoextra package in R Studio. In this analysis, variables with low Cohen’s F coefficient (effect size < 0.3) and outliers were discarded. The DNA sequences of the host molecular marker – mitochondrial - cytochrome C oxidase I gene (COI) were amplified using specific primers (Table S2). The PCR was performed in a reaction mixture consisting of 10 μl of the PCR Mix Plus mixture (A&A Biotechnology), 8 μl of water, 0,5 μl of each of the primers (10 μM) and 1 μl of the DNA template (1 μg/μl) under the following conditions: an initial denaturation step at 94°C for a duration of 3 min, followed by 33 cycles at 94°C for 30 s, 48-52°C for 40 s, 70°C for 1 min 40 s and a final extension step of 5 min at 72°C. The PCR products were made visible through electrophoresis in 1.5% agarose gel stained with Simply Safe (EURx) and then sequenced using the Sanger method (Genomed, Poland). All the nucleotide sequences obtained were deposited in the Figshare database under project “Typhlocybinae and Eurymelinae microbiomes”.

### Phylogenetic and cophylogenetic analyses

The obtained nucleotide sequences were aligned using ClustalW. Phylogenetic analyses were performed based on the 16S rRNA genes of *Sulcia* symbionts and COI genes of insects. The phylogenetic analyses were conducted using MrBayes software (86). For the Bayesian inference, four incrementally Metropolis coupling the MCMC chains (3 heated and 1 cold) were run for five million generations with trees sampled every 1000th generation. FigTree 1.3.1 and iTOL software were used for the visualization of the results of the Bayesian analysis.

### Fluorescence *IN SITU* hybridization (FISH)

Two methods of FISH were conducted in this work: whole-mount FISH and resin sections FISH. Whole-mount FISH technique by using tissue scanning allowed to make 3D models construction of bacteriomes and infected oocytes. Resin sections FISH technique allowed to visualize symbionts at high magnification, with better image quality. For the FISH method, the adult females preserved in 90% ethanol were rehydrated, fixed in 4% paraformaldehyde for two hours at room temperature and dehydrated through incubations in 80%, 90% and 100% ethanol and acetone. For whole-mount FISH, entire insects were also bleached in 6% hydrogen peroxide in 80% ethanol for two weeks to quench autofluorescence in the insect tissue. For resin sections FISH, the material was embedded in Technovit 8100 resin and cut into sections (1 µm thick) using an ultramicrotome Leica EMUC7. Hybridization was performed using a hybridization buffer containing: 1 ml 1M Tris-HCl (pH 8.0), 9 ml 5M NaCl, 25 μl 20% SDS, 15 ml 30% formamide and about 15 ml of distilled water. The slides or whole samples were incubated in 200 μl of hybridization solution (hybridization buffer + probes) overnight at room temperature (87). After this, the samples were washed in PBS three times for 10 minutes, dried and covered or suspended in a medium containing antifade reagent with DAPI fluorochrome (ProLong Gold, Life Technologies) then covered with cover glass. The hybridized samples were examined using a confocal laser scanning microscope Zeiss Axio Observer LSM 710 and ZEN (Zeiss) software. The fluorescence *in situ* hybridization experiments were repeated on three different individuals from each species (together 126 individuals). The probes and combined with them fluorochromes are presented in Table S2.

### Transmission electron microscopy (TEM)

Specimens collected in the field were dissected and preserved using 2.5% glutaraldehyde in 0.1 M phosphate buffer with pH = 7.4. The fixed material was stored at a temperature of 4 °C for a duration of three months, the specimens were then rinsed in the same buffer with an addition of sucrose (5.8 g per 100 ml) and postfixed in 1% osmium tetroxide for 1.5 h. Finally, after being dehydrated in a graded series of ethanol and acetone, the material was embedded in epoxy resin (Sigma Epoxy Embedding Medium kit). Using ultramicrotome Leica EMUC7 epoxy blocks were cut into sections. Semithin sections (1 µm thick), stained with 1% methylene blue in 1% borax, were used for the histological study (light microscope Nikon Eclipse E200). Ultrathin sections (70 nm thick), contrasted with uranyl acetate and lead citrate were used for the ultrastructural study under an electron transmission microscope Jeol JEM 2100 (80 kV) and EM-MENU 4 (TVIPS) software. The ultrastructural experiments were repeated on three different individuals from each species (together 126 individuals).

## Acknowledgment

This work was supported by the National Science Centre, Poland, Sonata 17, project no. 2021/43/D/NZ8/02183 to M.K.

## Author Contributions

Michał Kobiałka, Conceptualization, Data curation, Formal analysis, Funding acquisition, Investigation, Methodology, Project administration, Resources, Software, Supervision, Validation, Visualization, Writing – original draft, Writing – review and editing

Dariusz Świerczewski, Resources, Writing – review and editing

Marcin Walczak, Resources, Writing – review and editing

Weronika Urbańczyk, Data curation, Investigation

The authors declare no conflict of interest.

## Data Availability

Sequence data have been deposited in Figshare under project link: https://figshare.com/projects/Typhlocybinae_and_Eurymelinae_microbiomes/220621

## Supplementary Material

The following material is available online.

Table S1. All collected samples, including full names, abbreviations, place of collection, host plant, and the trophic specialty. Table S2. Used primers, including purpose, name, sequence, target gene, annealing temperature and source.

Fig. S1. The percentage relative abundance of bacteria for each Typhlocybinae sample based on the number of the sequence reads.

Mov. 1. *M. vicina,* the 3D view of a bacteriome consisted of *Sulcia* (green) bacteriocytes and the fragment of the fat body filled within *Ophiocordyceps* yeast-like symbionts (red).

Mov. 2. *P. albicans,* the 3D view of a bacteriome consisted of two zones, formed by bacteriocytes filled with *Sulcia* (green) and *Nasuia* (red) bacteria.

Mov. 3. *I. stigmaticalis,* the 3D view of a fragment of a bacteriome with bacteriocytes filled with *Sulcia* bacteria (green) infected by *Arsenophonus* (red).

Mov. 4 and 5. *P. albicans,* the 3D view of a "cap-like symbiont ball" consisted of *Sulcia* (green) and *Nasuia* (red) bacteria.

